# Anillin propels myosin-independent constriction of actin rings

**DOI:** 10.1101/2020.01.22.915256

**Authors:** Ondřej Kučera, Daniel Janda, Valerie Siahaan, Sietske H. Dijkstra, Eliška Pilátová, Eva Zatecka, Stefan Diez, Marcus Braun, Zdenek Lansky

**Affiliations:** Institute of Biotechnology of the Czech Academy of Sciences, BIOCEV, Vestec, Prague West, Czechia; B CUBE Center for Molecular Bioengineering, TU Dresden, Dresden, Germany; Max Planck Institute of Molecular Cell Biology and Genetics, Dresden, Germany; Cluster of Excellence Physics of Life, Technische Universität Dresden, Dresden, Germany

## Abstract

Constriction of the actin cytokinetic ring is an essential step of cell division. In a generally accepted view, the constriction is driven by relative sliding of actin filaments propelled by myosin motors. However, in multiple organisms, the ring constriction is myosin independent. How actin rings constrict in the absence of motor activity remains unclear. Here, we demonstrate that actin contractility can be propelled by anillin, a diffusible non-motor actin crosslinker, localising to the cytokinetic ring. We in vitro observed the formation and constriction of rings comprising multiple actin filaments bundled by anillin. Rings constricted due to anillin-generated forces maximising the overlap lengths between the filaments. Actin disassembly promoted constriction. We propose that actin crosslinkers, generating forces complementary to molecular motors, contribute to the contractility of diverse actin structures, including the cytokinetic ring.

## Introduction

Constriction of the cytokinetic actin contractile ring drives the division of most eukaryotic cells at the end of mitosis or meiosis (*1, 2*). The contractile ring is composed of bundles of actin filaments overlapping in mixed orientations, non-muscle myosin II motors, crosslinking proteins and scaffolding proteins (*3*–*5*). It was suggested that the ring constriction is driven by myosin-propelled relative sliding of actin filaments, analogous to muscle sarcomere contraction (*6, 7*). Unlike muscle sarcomeres, however, the orientation of actin filaments forming the ring is disordered (*8*). The sliding activity of myosin alone in this disordered system is thus equally likely to locally lead to contraction or extension and is insufficient to generate any net constriction of the ring (*9*). Additional factors are therefore required to locally break the symmetry of the system in order to favour contractile forces (*3, 9*–*11*). Strikingly, in multiple organisms, myosin-II activity or presence is not required for cytokinesis. In budding yeast, the rings constrict when myosin-II motor activity is disabled (*12*), budding yeast and *Dictyostelium* cells constrict when myosin-II is deleted (*13, 14*), and a myosin-II-independent phase of the ring closure was identified during *Drosophila* embryo cleavage (*15*). These findings demonstrate that distinct mechanisms, myosin-independent and myosin-dependent, drive the constriction of cytokinetic actin rings.

Theoretical work and *in vivo* experiments suggest that possible sources of the driving force underlying myosin-independent constriction mechanisms depend on actin-crosslinking proteins and actin filament disassembly (*16*–*20*). In microtubule networks, similar force-generating mechanisms, independent of molecular motors, are directly experimentally observed. Confinement of diffusible crosslinkers in regions of overlap between microtubules, resulting in the generation of entropic forces, leads to directional microtubule-microtubule sliding (*21, 22, 24*). Kinetochores couple to the disassembling microtubule tips (*23*) to drive the movement of chromosomes. Direct experimental evidence of crosslinker-propelled actin filament sliding is, however, missing, leaving the question of the myosin-independent mechanism of actin ring constriction unanswered.

Here we show in a minimal reconstituted system that constriction of actin rings can be propelled by anillin, non-motor actin crosslinking and scaffolding protein highly enriched in the contractile cytokinetic ring during cytokinesis (*25*–*27*). We find that anillin directly drives the relative sliding of actin filaments and can couple with actin filament disassembly to generate contractility. We thus demonstrate that filament crosslinkers, such as anillin, can generate contractile forces in actin networks, substituting for molecular motor activity.

## Results

### Anillin slides actin filaments to maximize their overlap

To study the interactions between anillin and actin filaments *in vitro*, we specifically immobilised sparsely rhodamine-labelled, phalloidin-stabilised actin filaments to the coverslip surface (Methods). After the addition of GFP-labelled anillin to the experimental chamber, using TIRF microscopy, we observed anillin-GFP binding to actin filaments (Fig. S1A). At the anillin-GFP concentration of 0.12 nM, actin filaments were decorated by anillin-GFP molecules (Fig. 1 A, B), which in accordance with previously published data (*26*) we identified as monomers (Fig. S1 B). These single anillin-GFP molecules diffused along the actin filaments with a diffusion constant of 0.0088 ± 0.0006 μm^2^s^-1^ (linear fit coefficient ± 95% confidence bounds, 268 molecules in 2 experiments) (Fig. S1 C). When we increased the anillin-GFP concentration in solution to 12 nM and simultaneously added brightly rhodamine-labelled, phalloidin-stabilised actin filaments (mobile filaments), we observed mobile filaments landing from solution and length-wise crosslinking with the immobilised filaments forming filament bundles. In these bundles, we found mobile filaments moving diffusively along the immobilised filaments (Fig. 1 C, Movie S1), showing that anillin-GFP generates a diffusible link between actin filaments. Strikingly, when a mobile filament landed such that it overlapped partially with the immobilised filament, we observed these mobile filaments moving directionally along the immobilised filaments. Importantly, this movement was always in the direction increasing the length of the overlap between the two filaments and thus effectively contracting the elementary two-filament actin bundle (Fig. 1 D, Movie S2). We performed this experiment also in an alternative way, not immobilising any of the filaments, but imaging the bundle formation in solution in the presence of 0.1% (w/v) methylcellulose to facilitate the imaging at the plane near the coverslip surface (Methods). Also, in this set of experiments, we observed, upon an initial formation of a partially overlapping two-filament bundle, relative directional sliding of the two crosslinked filaments increasing their overlap length (Fig. 1 E, Movie S3). The presence of anillin was required for the formation of the actin bundles and their directional sliding (Fig. S1 D, Movie S4). We note that in both types of experiments, we neither observed directional sliding of filaments that overlapped fully nor directional sliding decreasing the overlap length. We also note that the directional sliding did not depend on the relative orientation (structural polarity) of the bundled filaments (Fig. S1 E, F). Analysing the dynamics of the sliding, we found that the overlap length started increasing immediately after the formation of the bundle. Concomitantly with the increase of the overlap length, anillin-GFP amount in the overlap increased (Fig. S1 G), indicating anillin-GFP binding into the newly forming overlap region. The sliding was slowing down as the overlap length, and the amount of anillin in the overlap increased, leading to the overlap length reaching an equilibrium value (Fig. 1 F, G, Fig. S1 H, I). The mean difference between the initial and final overlap length, i.e. the mean contraction length, was 1.20 ± 0.62 μm (± s.d., maximum 2.67 μm, n = 24 actin filament pairs). Combined, these data demonstrate that anillin drives directional sliding of actin filaments relative to each other, increasing the length of the overlap between the filaments and thus contracting the filament bundle.

**Fig. 1.**
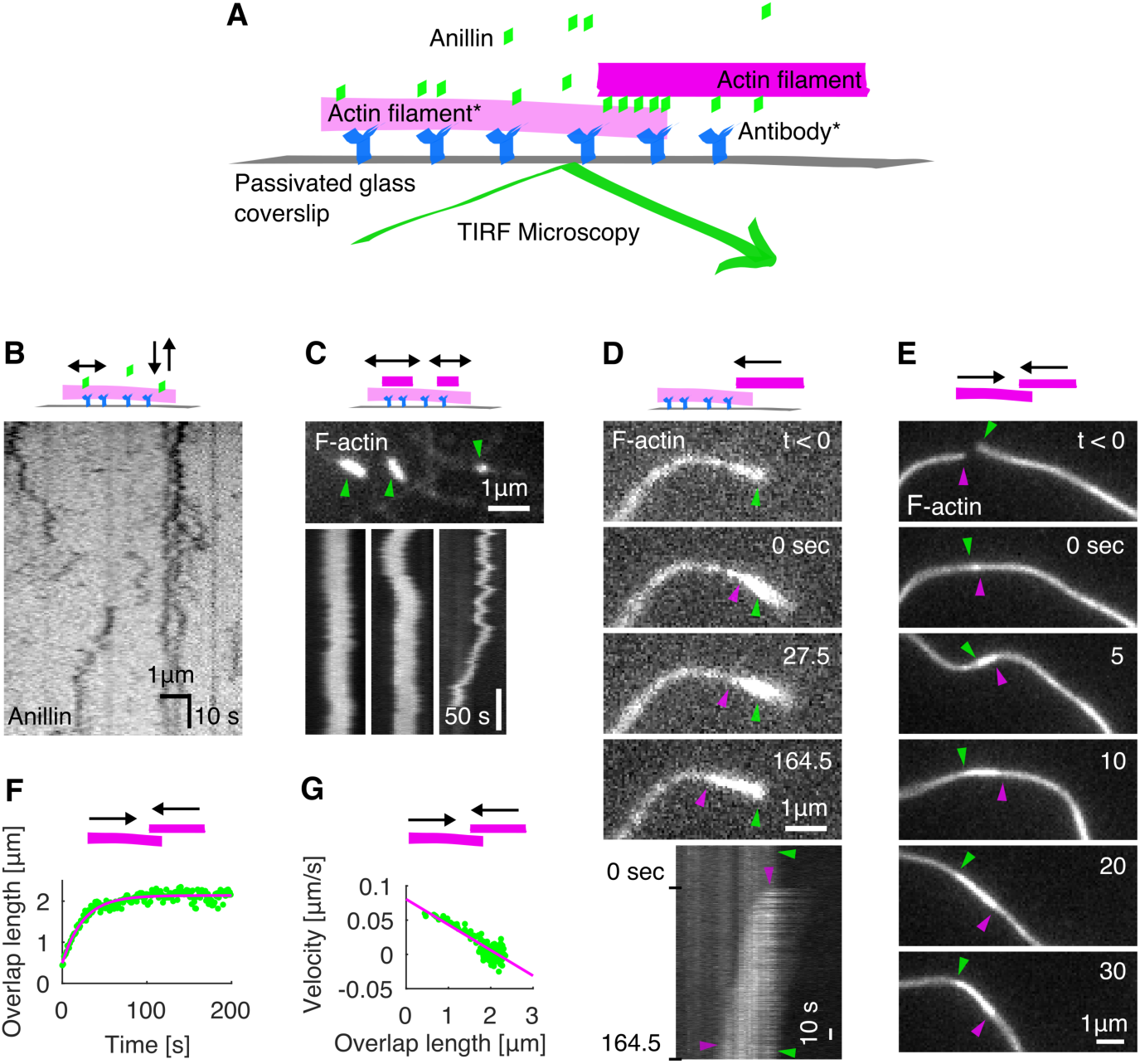
Anillin slides actin filaments to maximize their overlap. (**A**) Schematic representation of the experimental setup. The asterisk denotes components used only in experiments with immobilised filaments. (**B**) Intensity-inverted kymograph showing single anillin-GFP molecules diffusing along an actin filament. (**C**) Fluorescence micrograph (top) showing three short mobile actin filaments (indicated by green arrowheads) crosslinked by anillin-GFP (not visualized) to long, sparsely fluorescently labelled, immobilised actin filaments (see Movie S1). Kymographs (bottom) show the diffusion of mobile actin filaments from the micrograph above along the immobilised filaments. (**D**) Time-lapse micrographs (top) and a kymograph (bottom) showing anillin-driven sliding of a mobile actin filament (bright) along an immobilised filament (dim), increasing the overlap between the two actin filaments (see Movie S2). Green and magenta arrowheads indicate the ends of the immobilised and mobile filaments, respectively. This experiment was repeated 5 times (8 events observed) with similar results. (**E**) Time-lapse micrographs showing anillin-driven sliding of two mobile actin filaments along each other, increasing the overlap between the two actin filaments (see Movie S3). Arrowheads indicate the filaments’ ends. This experiment was repeated 15 times (24 events observed) with similar results. (**F**) A typical time trace of the overlap expansion (green dots) reaching an equilibrium value (magenta line represents an exponential fit (Methods) to the data). (**G**) Velocity of the overlap expansion decreases with increasing overlap length. Green points represent an exemplary event, magenta line is a linear fit to the data.

### Anillin couples with actin disassembly to generate directed filament sliding

As actin depolymerization is an important factor for the constriction of the cytokinetic ring, we wondered how will depolymerization of actin filaments affect the observed anillin-driven filament sliding. We employed the above described experimental setup using a mixture of rhodamine-labelled, phalloidin-stabilized actin filaments and unlabelled non-stabilized actin filaments in the presence of 12 nM anillin-GFP and not immobilising any of the filaments. To enhance the filament disassembly, we used 80-200 nM latrunculin (*28*). Similar to the experiment with stabilized actin, we observed anillin-dependent crosslinking of the filaments leading to the formation of actin bundles, which contracted over time. Strikingly, however, in this experiment, we did not observe only sliding of partially overlapping filaments as described above, but we also observed sliding of filaments that overlapped fully. This sliding was observed when the retreating tip of the disassembling non-stabilized actin filament reached a stabilized filament (Fig. 2 A, Movie S5). The stabilized filament then followed the retreating tip such that the velocity of sliding matched the rate of the filament disassembly (correlation coefficient 0.90 ± 0.06, mean ± s.d., n = 13 events in 9 experiments) (Figs. 2 B-D) and thus a full overlap between the two filaments was maintained (Fig. 2 A). During this sliding, the Anillin-GFP density in the overlaps increased with a faster rate than on single filaments (Fig. S2), suggesting that the overlap ends form boundaries for anillin diffusion. As the disassembly of one of the filaments works towards shortening the overlap length, the concurrent anillin-driven sliding compensates this shortening, maintaining the overlap length. This process, leading to the sliding of actin filaments along with the tip of a depolymerizing filament, is thus an efficient mechanism that couples actin filament depolymerization into sliding-driven contraction of actin bundles.

**Fig. 2.**
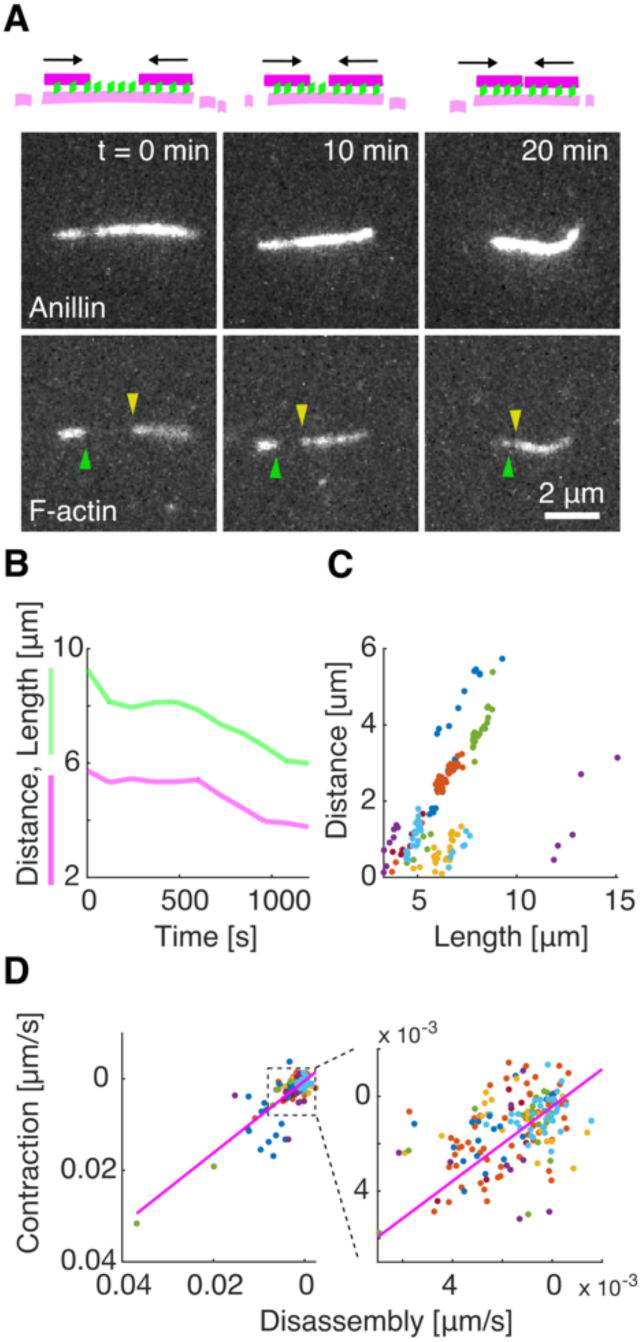
Anillin couples with actin disassembly to generate directed filament sliding. (**A**) Fluorescence time-lapse micrographs showing two short phalloidin-stabilised filaments sliding along with the retreating ends of a non-stabilised filament disassembling in the presence of latrunculin (see Movie S5). The disassembling filament is not visualized directly, its length is visualized by the anillin-GFP signal. Arrowheads indicate the ends of the two stabilized actin filaments. (**B**) Typical time-traces of the length of the non-stabilised filament (green) and the distance between the two short stabilised filaments moving with the ends of the non-stabilised one. (**C**) The distance between the stabilized filaments decreases linearly with the length of the disassembling filament (n = 12 events in 9 experiments, colours denote individual events). (**D**) The contraction rate of the actin bundle scales linearly with the rate of disassembly of the non-stabilized filament (the same experimental data as shown in (C)).

### Anillin promotes formation of actin rings and drives their constriction

To study the anillin-driven actin filament sliding in structures of higher complexity, we increased the actin filament density in the solution, not immobilizing any of the filaments. Initially, we used non-stabilised actin filaments. We found that anillin, similar as septins (*29*), promotes the formation of rings of actin filaments several micrometres in diameter (Figs. 3 A,B, S3), which we observed forming throughout the volume of the microscopy chamber. The rings typically consisted of 3 - 8 actin filaments colocalising with anillin-GFP (Methods). Importantly, time-lapse imaging of these rings revealed that they constrict over time (Fig. 3 B,C, Movie S6). The circumference of the rings decreased exponentially (Fig. 3 C-E), enabling us to quantify the asymptotical circumference of the maximally constricted ring. The constriction varied substantially, with maximum constriction after 300 s reaching 20 % of the initial circumference, maximum asymptotical constriction reaching 100 % and maximum constriction rate reaching 0.034 μm×s^-1^ (Fig. 3 F, G). The spread in the data likely reflects the uncertainty of the initial lengths of the overlaps between the filaments forming the rings, as well as the spread in the number of these filaments. While most of the rings (n = 18, out of total 26 rings in 16 experiments) constricted during the time of the experiment, in some cases (n = 8), we did not observe any significant constriction. We interpret these as rings that already reached its maximal constriction before the beginning of our observation. To test the contribution of actin disassembly in the constriction process, we studied the constriction dynamics of anillin-actin rings consisting of filaments stabilised by rhodamine-phalloidin. We found that the constriction was significantly slower (P = 0.0044), with maximum constriction after 300 s of about 5 % of the initial circumference and maximum rate of 0.006 μm×s^-1^ (Fig. 3 F, G, n = 12 rings in 10 experiments). However, the maximum asymptotical constriction reached 100 %, similarly to the rings formed by disassembling actin. By contrast, the addition of latrunculin to rings of non-stabilised actin filaments led to higher asymptotic constriction (Fig. 3 G). These results are in agreement with our data on linear bundles, where we observed that depolymerisation of actin filaments promotes the contraction of the bundle. Combined these experiments show that rings of actin filaments can constrict without the action of nucleotide hydrolysing molecular motors, solely driven by forces generated by passive actin crosslinkers. This constriction is enhanced by the actin filament disassembly.

**Fig. 3.**
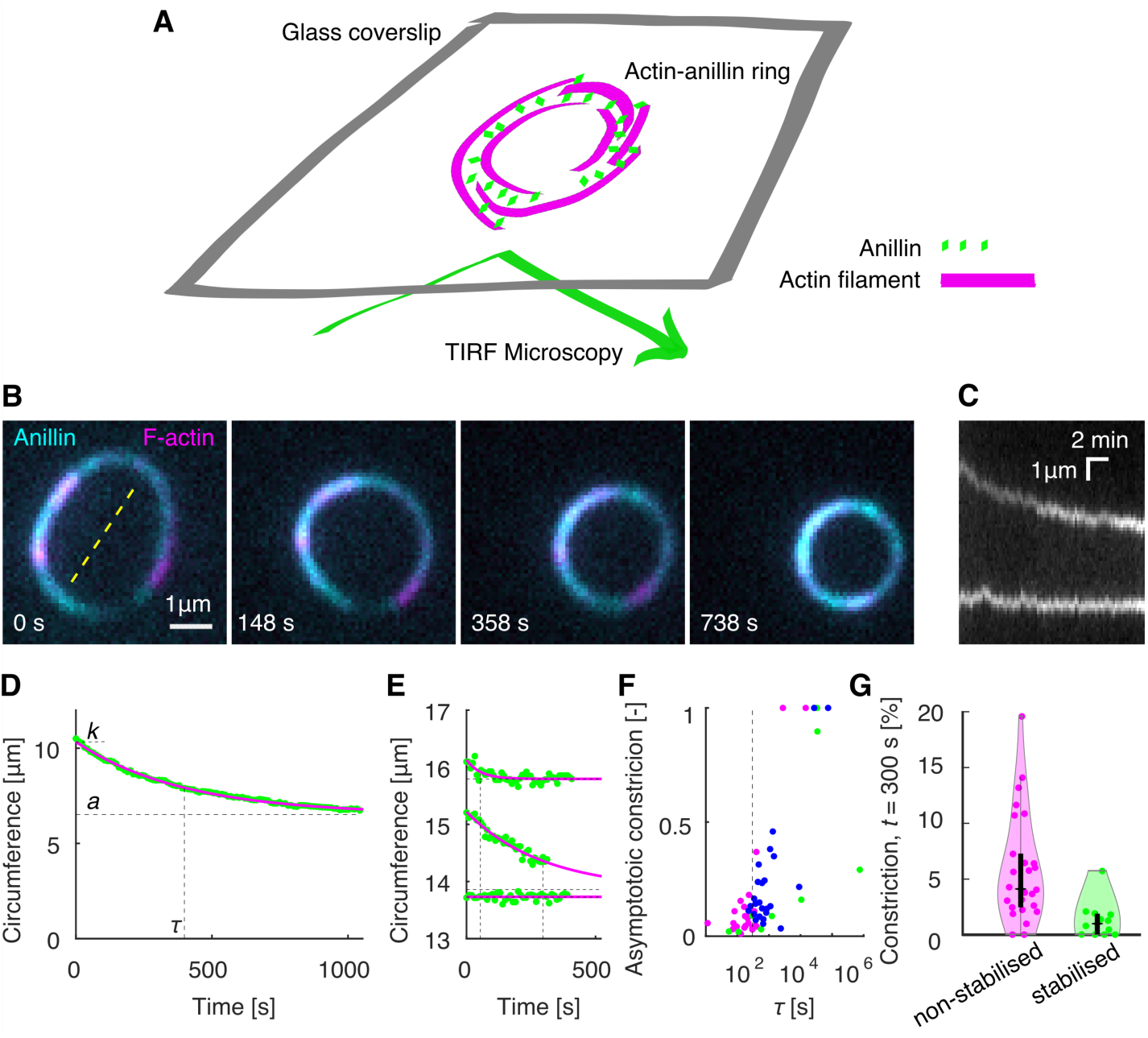
Anillin promotes formation of actin rings and drives their constriction. (**A**) Schematic representation of the assay geometry. (**B**) Multicolour time-lapse micrographs of the anillin-actin ring, showing the ring constriction over time (see Movie S6). (**C**) Kymograph through the yellow dashed line in (B) of the constricting anillin-actin ring. (**D**) The time trace of the circumference of the ring shown in (B) and (C). Green dots represent experimental data, and the solid magenta curve an exponential fit. Dashed lines indicate the fit parameters (*τ* – time constant, *k* – initial circumference, *a* – the asymptotical value of the circumference). (**E**) Exemplary time-traces of a fast (top), slow (centre), and not (bottom) constricting rings. Identical graphical representation as in (D). (**F**) Relation between the time constant and the relative asymptotic constriction (1 – (*a/k*)). Each dot represents one ring formed either by non-stabilised (magenta, n = 26 in 16 experiments) or stabilised (green, n = 12 in 10 experiments) actin, or non-stabilized actin in the presence of latrunculin (blue, n = 25 in 7 experiments). Fits leading to asymptotical constriction higher than 1 were cut off to 1 (n = 5). Events with higher time constant than 10^6^ s, all having zero asymptotic constriction, are not shown (n = 5). *τ* = 300 s is highlighted by vertical dashed line. (**G**) Percentage of the constriction at *t* = 300 s for all the rings of stabilised and non-stabilised actin (n = 38). Individual data points are accompanied with violin plots, and black boxplots (horizontal tick – median, box – interquartile ranges, whiskers – 95% confidence intervals).

### Anillin generates tens of pico-Newton forces to slide actin filaments

To directly measure the forces generated by anillin in the actin bundles, we specifically attached actin filaments to two microspheres (Methods). Holding the microspheres by two traps in an optical tweezers setup, we formed between them, in the presence of 12 nM anillin-GFP, a crosslinked actin bundle consisting on average of about 5 filaments (4.7 ± 1.3, mean ± s.d., n = 35 bundles) (Fig. 4 A, B, Methods, Fig. S4 A). We then moved the microspheres apart from each other in 100 nm steps, stretching the bundle and thus sliding the filaments within the bundle in the direction shortening their overlaps (Fig. 4B). Simultaneously with each step-movement of the trap (each step taking approx. 33 μs), we observed a rapid increase in force, which then decayed within several seconds to a plateau reflecting the equilibrium between the anillin-generated force and the external load exerted by the optical trap (Fig. 4 C, D, see Fig. S4B for control experiment). Since the median time constant of the force decay (1.06 s, quartiles 0.47s and 2.00 s, n = 326 steps in 43 experiments) together with the trap movement was an order of magnitude faster than the rate of anillin unbinding from actin (Fig. S1A, Methods), we interpret this dynamic response being due to the rearrangement of anillin molecules in the overlap. The equilibrium force values increased with increasing distance between the two microspheres and thus with decreasing overlaps between the filaments in the bundle, reaching up to tens of pico-Newtons, up to a point when the bundle ruptured (n = 326 steps in 43 experiments, Fig. 4 E, F, Fig. S4 C). When we (before the rupture) decreased the external load on the bundle moving the microspheres closer together in steps of 100 nm, in a situation when the bundle was pre-stretched and the force equilibrium was non-zero, we immediately observed relative movement of the two microspheres in the direction increasing the overlap length (Fig. 4 C, D) manifesting relative sliding of actin filaments as a direct consequence of the anillin-generated force. To test how the generated force depends on changes in the anillin concentration, we first pre-stretched the actin bundle by moving the microspheres apart such that the equilibrium force readout was non-zero, and then we decreased the concentration of anillin-GFP from 12 nM to 1 nM (keeping the distance between the microspheres and thus the external load constant, Methods). We then observed the force generated by anillin decreasing over time, presumably as anillin-GFP was unbinding from the overlap (Fig. 4 G, left panel, see Fig. S4D for control experiment). Conversely, increasing the anillin-GFP concentration in the measurement chamber resulted in a gradual increase in the anillin-generated force as anillin-GFP was binding into the overlap (Fig. 4 G, right panel). Combined, these results show that anillin generates substantial forces in the range of tens of pico-Newtons per bundle of about five filaments. These forces result in relative actin sliding increasing the overlaps between actin filaments in the bundle and, consequently, in the bundle contraction.

**Fig. 4.**
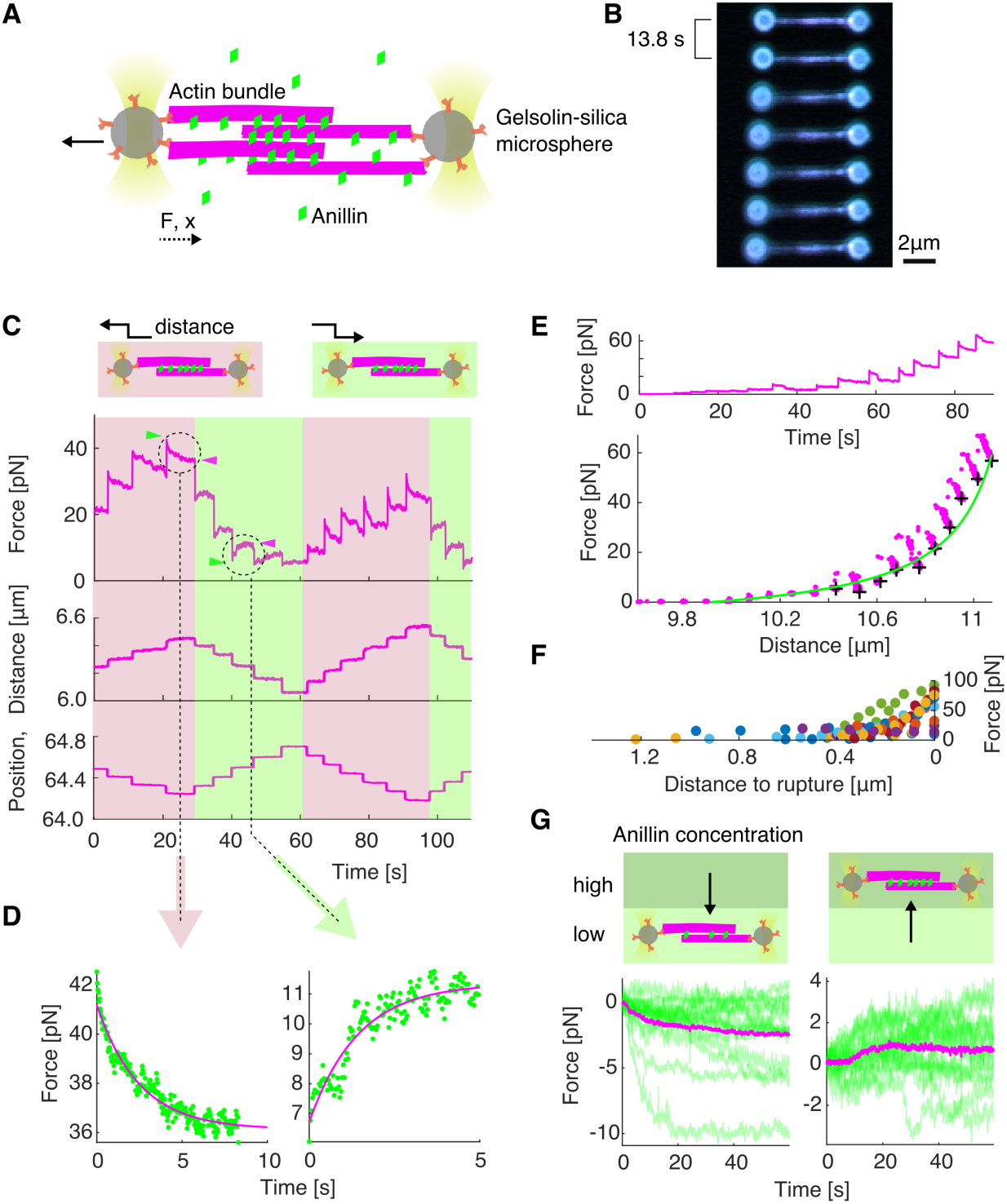
Anillin generates tens of pico-Newton forces to slide actin filaments. (**A**) Schematic representation of the experimental setup. (**B**) Multicolour time-lapse fluorescence micrographs showing an actin bundle attached between two silica microspheres (actin - green, anillin - blue). The bundle is being stretched as the left microsphere is pulled leftwards by an optical trap. (**C**) Typical temporal profile of the detected force (top) and the distance between the microspheres (middle) in response to the controlled step-wise movement of the left microsphere (bottom). A typical force response after stretching and relaxing steps are indicated by the light brown and light green background, respectively. The green arrowheads point to the initial peak force and the magenta arrowheads show the plateau force in exemplary steps. (**D**) Magnification of the two typical examples highlighted in (C) of the force response after a stretching (left) and relaxation (right) steps. Green dots represent individual time points, magenta curve is an exponential fit. (**E**) Typical force time-trace (top) and the force-distance curve (bottom) corresponding to stretching of an anillin-actin filament bundle (experimental data points - magenta). Asymptotic forces of individual stretching steps, calculated by fitting an exponential to the force decays (as shown in (D)), are indicated by black crosses. These increase hyperbolically with increasing distance between the microspheres, and thus with decreasing overlap length *L*. Green line represents ∼1/*L* fit to the data. (**F**) Detected force increases with the decreasing distance before a rupture of the bundle - with decreasing overlap length. All events longer than 6 steps are plotted (n = 11 experiments indicated by different colours). (**G**) Force response of a pre-stretched actin-anillin bundle to a change of anillin-GFP concentration. Schematic representation of the experiment (top) and temporal experimental data (bottom). Left: decrease of the concentration (n = 15 events in 14 experiments), right: increase of the concentration (n = 15 events in 14 experiments). Green curves are the experimental data, mean temporal profile is shown in magenta.

## Discussion

We here show that passive, non-motor, actin crosslinker anillin can drive the contraction of actin bundles through directed filament sliding. Depletion interactions (*30*–*33*) did not contribute to this phenomenon, as filaments did neither bundle nor slide in the absence of anillin in our control experiments (Fig. S1 D). Anillin creates diffusible links between actin filaments (Fig. 1 B, C) and is retained in overlaps that transiently shorten (Fig. S2), suggesting that anillin molecules in the overlap can be described as confined gas, analogous to crosslinkers described in microtubule systems (*21, 22*). We observed that anillin generated force increased hyperbolically with decreasing overlap length (Fig. 4 E, F) and, as anillin unbinding on the experimental timescale was negligible, likely thus with increasing density of anillin in the actin overlap, as expected for an ideal gas (*22*). Increasing or decreasing the concentration of anillin in our trapping experiment resulting in increasing or decreasing anillin-generated force, can be interpreted as an increase or decrease in the anillin-gas pressure in the overlap. Combined, thus, our results suggest that the anillin-generated force is of an entropic origin (*34*) with an additional force component likely associated with energetically favourable binding of the anillin crosslinkers into the overlap, analogous to microtubule crosslinkers (*21, 22*). Anillin crosslinkers likely induce friction between the filaments that they bundle (*22, 35*), which might explain the slowdown in sliding or ring constriction observed with increasing overlap lengths. Circular constraints within the ring may also contribute to the deceleration of the constriction as the decrease of the diameter of the ring requires bending of its constituting filaments (*29, 33, 36*). At anillin concentrations of 12 nM used in this study, anillin molecules, collectively, can generate forces in the order of 10 pN in a bundle of about five filaments. These forces are comparable to forces generated by myosins; single Myosin-II stall force is about 2 pN (*37, 38*), suggesting that anillin crosslinkers can generate forces relevant in the context of the cytokinetic ring.

In a disordered actin array, such as in the cytokinetic ring, myosin motors alone are equally likely to locally generate contractile or extensile forces. Additional mechanisms are needed to break the symmetry of the process, locally favouring the contractile forces to drive the overall net constriction of the ring. In this context, various factors are currently under debate (reviewed extensively in (*3, 10, 11*)), such as the difference in behaviour of actin filaments under compressive and extensile forces (*39, 40*), inhomogeneous distribution of myosin motors (*18, 41*–*43*) or membrane anchoring of actin filaments (*38*). Importantly, connectivity between actin filaments mediated by actin crosslinker *α*-actinin facilitates myosin-driven contraction of actin rings in vitro (*9*). We here show that anillin-generated forces always act in the direction increasing the overlap length, contracting the actin bundle. Anillin will thus locally generate assisting forces in overlaps, which are locally contracted by myosin and resisting forces in overlaps, which myosin locally extends. Therefore, anillin generated force could be an element breaking the symmetry for myosin-dependent force generation in the cytokinetic ring and thus, in concert with myosin, drive its constriction. This notion is consistent with *in vivo* results showing that, in *Drosophila* embryos, lack of anillin results in a complete stall of myosin-driven contraction in the myosin-dependent phase of the ring constriction (*15*).

Actin filament depolymerization in various, albeit not all, organisms (*44*) plays an important role during the constriction of the cytokinetic ring. Blocking actin depolymerization, e.g. with drugs or by impairing the actin-depolymerizing and severing enzyme cofilin, leads to defective ring constriction (*20, 45*–*47*). Moreover, decreased levels of cofilin or anillin resulted in similar effect in *Drosophila* embryos, namely a delayed switch to the myosin-independent phase of the constriction (*15*), suggesting that actin filament depolymerization, in concert with anillin might drive myosin-independent constriction of the cytokinetic ring. We here provide direct evidence that anillin crosslinkers efficiently couple with actin filament disassembly to generate contractile forces. This mechanism is reminiscent of the experimentally observed mechanism driving chromosome movement during the anaphase of cell division by coupling the chromosomes’ kinetochores to depolymerizing microtubule ends by biased diffusion (*23, 48, 49*) as proposed earlier theoretically (*50*). Analogously, the diffusion of anillin is likely biased away from depolymerizing actin filament ends, resulting in directional sliding of crosslinked filaments with the depolymerizing end. We hypothesize that severing of actin filaments might promote this process, by increasing the number of filament ends, which can trigger at these positions additional anillin-mediated filament sliding and accelerate the bundle contraction.

In summary, we show directly that actin crosslinkers can drive the contraction of actin bundles and can couple with actin disassembly to enhance this effect. We propose that this mechanism can i) underpin the *in vivo* observed myosin-independent constriction of cytokinetic rings and ii) break the symmetry of the randomly oriented actin-myosin system in the ring enabling myosin-dependent constriction of the ring. A recent finding that anillin promotes tensile forces in the apical actomyosin networks of *X. laevis* embryonal epithelium (*51*) suggests that analogous mechanism might also be employed in other actin structures. We propose that crosslinker-dependent contractility of filamentous networks is a fundamental mechanism readily available in actin-based cytoskeletal structures, likely contributing to various cellular movements.

## Supporting information

S1_Diffusion_filaments

S2_SlidingON_AB

S3_SlidingON_methylcellulose

S4_Control_no_anillin

S5_Disassembly_contraction

S6_Ring

## Acknowledgments

We thank Andrew Ward for hints on actin crosslinking in the optical tweezers, the Protein Facility of MPI-CBG and Yulia Bobrova for technical support and Laurent Blanchoin for critical reading of the manuscript. This study was supported by the projects nos. 19-27477X to Z.L. and 20-04068S to M.B. from the Czech Science Foundation, project Introduction of New Research Methods to BIOCEV (CZ.1.05/2.1.00/19.0390) from the European Regional Development Fund, and by the institutional support of the Czech Academy of Sciences (RVO: 86652036). We acknowledge Biophysics facility of CMS supported by MEYS CR (LM2015043) and the Centre of Imaging Methods core facility, Faculty of Science, Charles University, supported by the Czech-BioImaging through MEYS CR (LM2015062 and CZ.02.1.01/0.0/0.0/16_013/0001775). DJ and EP carried out this research partly as a project work at the Faculty of Biomedical Engineering of the Czech Technical University in Prague.

## Author contributions

Conceptualization, MB, ZL; Methodology, OK, MB, ZL; Investigation, OK, DJ, VS, SHD, EP; Formal Analysis, OK, DJ, EP; Data curation, OK, DJ, SHD; Validation, OK; Resources, OK, EZ, SD; Writing, OK, MB, ZL; Visualization, OK; Supervision, MB, ZL; Funding acquisition, MB, ZL.

---

**Fig. S1.**
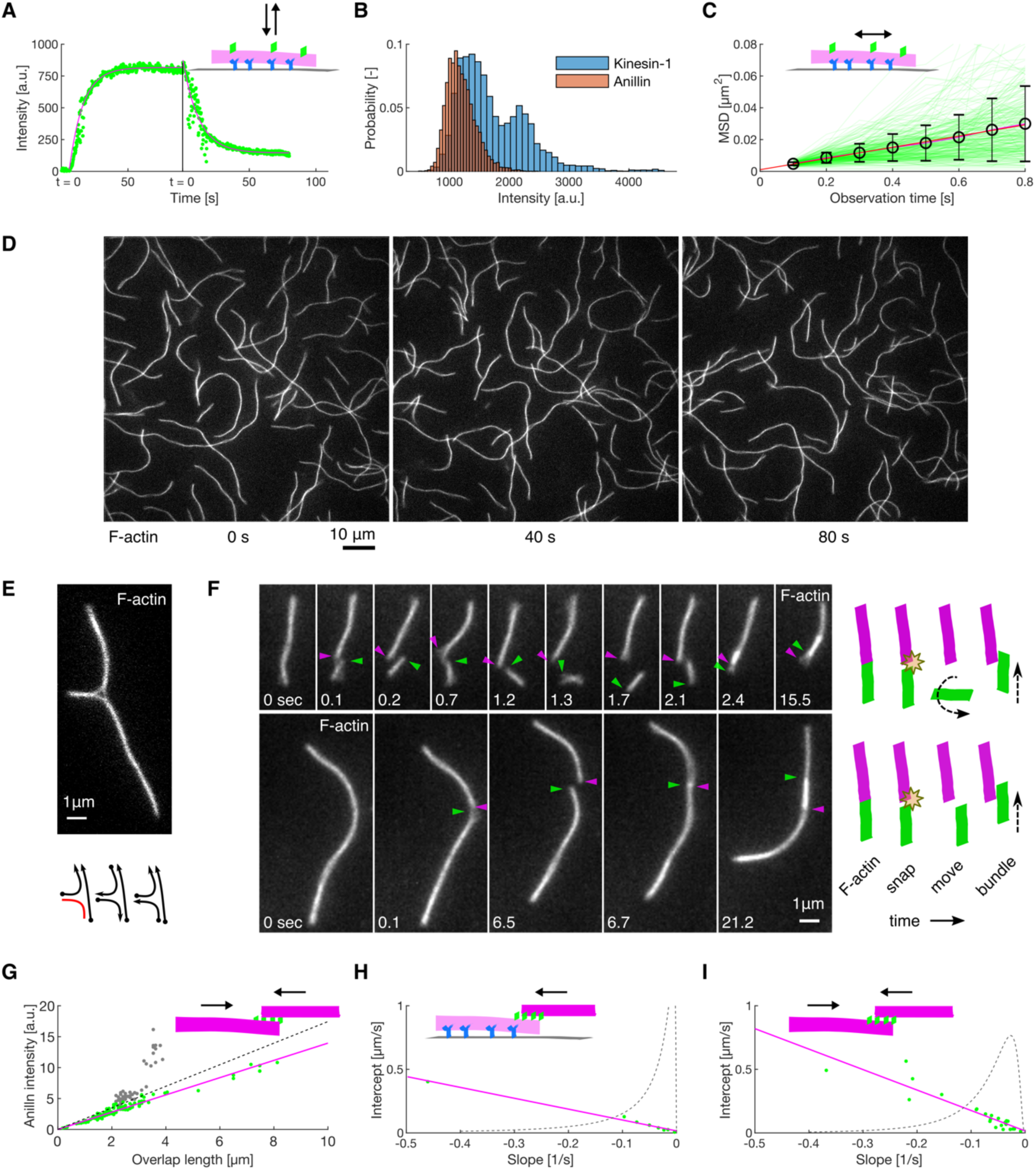
(**A**) Kinetics of the anillin-GFP binding/unbinding to/from actin filaments. A temporal profile of the fluorescence signal of the anillin-GFP along an actin filament shows a typical first-order growth upon addition of 6 nM anillin-GFP and first-order decay after the washout. The loading experiment was repeated 15 times (3 experiments) with similar result. The estimated unbinding rate of anillin from actin filaments is *k*off = 0.06 s^-1^, 95% confidence interval 0.03 – 0.10 s^-1^, 15 measurements in 3 experiments. (**B**) Histograms of the fluorescence intensity of single dimeric GFP-labelled kinesin-1 molecules bound to a microtubule (blue) and anillin-GFP particles bound to an actin filament (red). The histogram of kinesin-1-GFP shows typical bimodal shape, while that of anillin-GFP is mono-modal suggesting that anillin is in monomeric state (128 anillin-GFP molecules in 2 replicates). Note that the fluorescence intensities in (A) and (B) are not comparable as they were recorded with different acquisition times and using different excitation laser powers. (**C**) Mean-squared displacement (MSD) of anillin-GFP molecules diffusing on actin filaments. Green curves show MSD traces of individual molecules (n = 268 molecules in 2 experiments), the error bars show mean ± SD. Red line represents a linear fit to the mean of the MSD. (**D**) Actin filaments do not bundle in the absence of anillin (see Movie S4). Time-lapse micrograph shows that actin filaments are free to move in the lateral plane, but do not form bundles when anillin is absent, showing that the concentration of methylcellulose used in experiments with no immobilised filaments is low enough to not induce actin bundling through depletion effects (*8*). (**E, F**) Anillin can bundle and slide both parallel and antiparallel actin filaments. (E) A construct of three filaments bundled by anillin-GFP in which every filament overlaps partially with each other. Below is a schematic illustration showing that both, parallel and antiparallel filament pairs are formed, in agreement with previously published data (*9*). (F) Time-lapse fluorescence micrographs showing initially a single actin filament in the presence of anillin (not visualized). After the filament broke due to thermal fluctuations, the two parts connected again in an antiparallel (top) and parallel (bottom) manner, respectively, and started sliding along each other. The arrowheads indicate the newly formed filament ends formed by the breaking of the initial filament. A schematic representation of the events is shown to the right. (**G**) Intensity of anillin-GFP in the overlap during relative sliding of unbound actin filaments. The amount of anillin-GFP in the overlap between two filaments scales linearly with the overlap length. The dots represent individual timepoints (n = 187 time points in 16 events, 5 experiments). The data were fitted by a linear trend (dashed line). Two events with unusually higher intensity were marked as outliers (grey dots). The linear trend of remaining events (green dots, n = 142 time points in 14 events, 4 experiments) is shown in solid line (magenta). (**H, I**) Dependence between the parameters of the linear fit to the data, such as shown in Fig. 1 G (sliding velocity as a function of the overlap length) in experiments using (H) immobilised actin filaments (n = 8 bundles in 5 experiments) and (I) mobile actin filaments in the presence of methylcellulose (n = 24 bundles in 15 experiments). Although the events had random initial overlap length, the contraction in all events followed a regular pattern with linear dependence between the extrapolated maximum sliding velocity (intercept) and the deceleration rate (slope). Green dots show data from individual sliding events, solid magenta line is a linear trend, and the dotted curve shows an estimate of the exponential distribution of the slope of the fit.

**Fig. S2.**
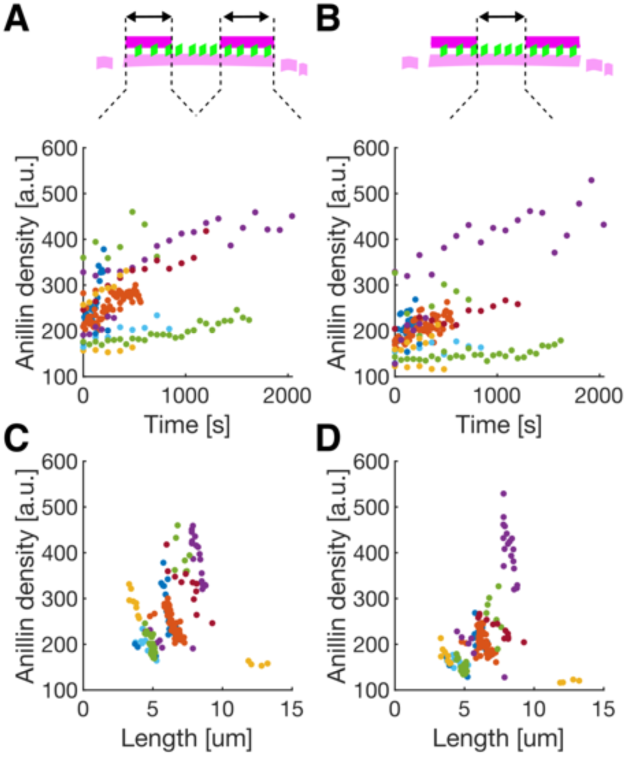
The density of anillin (mean fluorescence intensity of anillin-GFP per unit length of the actin filament) as a function of time, in the overlaps **(A)**, and along the single filament between the two overlaps **(B)** during the disassembling experiments (as shown in Fig. 2 A) and **(C**,**D)** as a function of the length of the bundle. Individual events are colour-coded (n = 181 time points in 12 events, 9 experiments). As the bundle contracts, the density of anillin is increasing on average 1.39 times (95% confidence interval 1.34 – 1.46; fit to the temporal profiles when normalised to the initial density) faster in the overlaps compared to single filament.

**Fig. S3.**
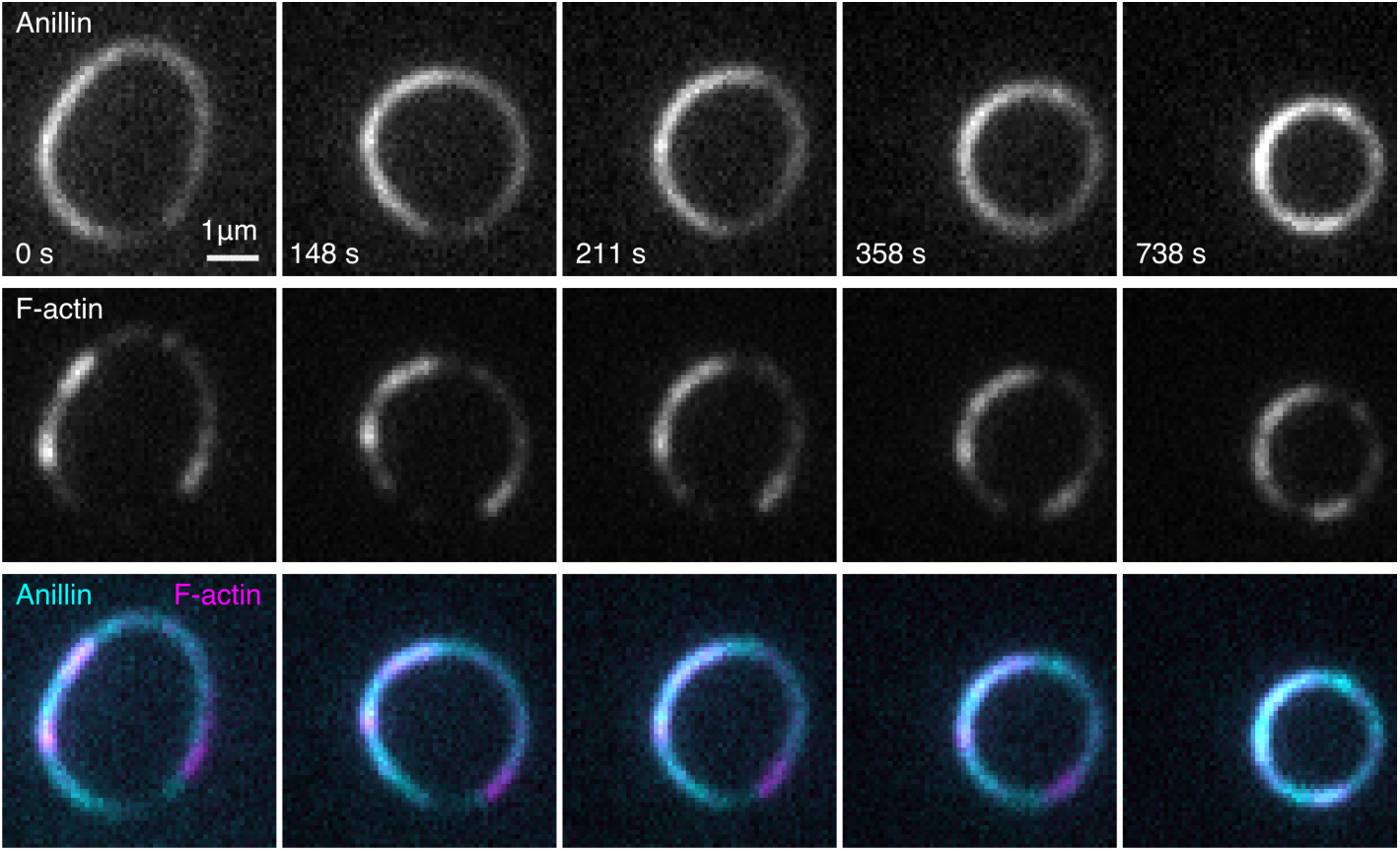
Time-lapse micrograph of the constriction of the anillin-actin ring composed of non-stabilised actin filaments, showing the anillin-GFP and rhodamine-actin channels and their overlay (see Movie S5). The experiment was repeated 16 times with similar results.

**Fig. S4.**
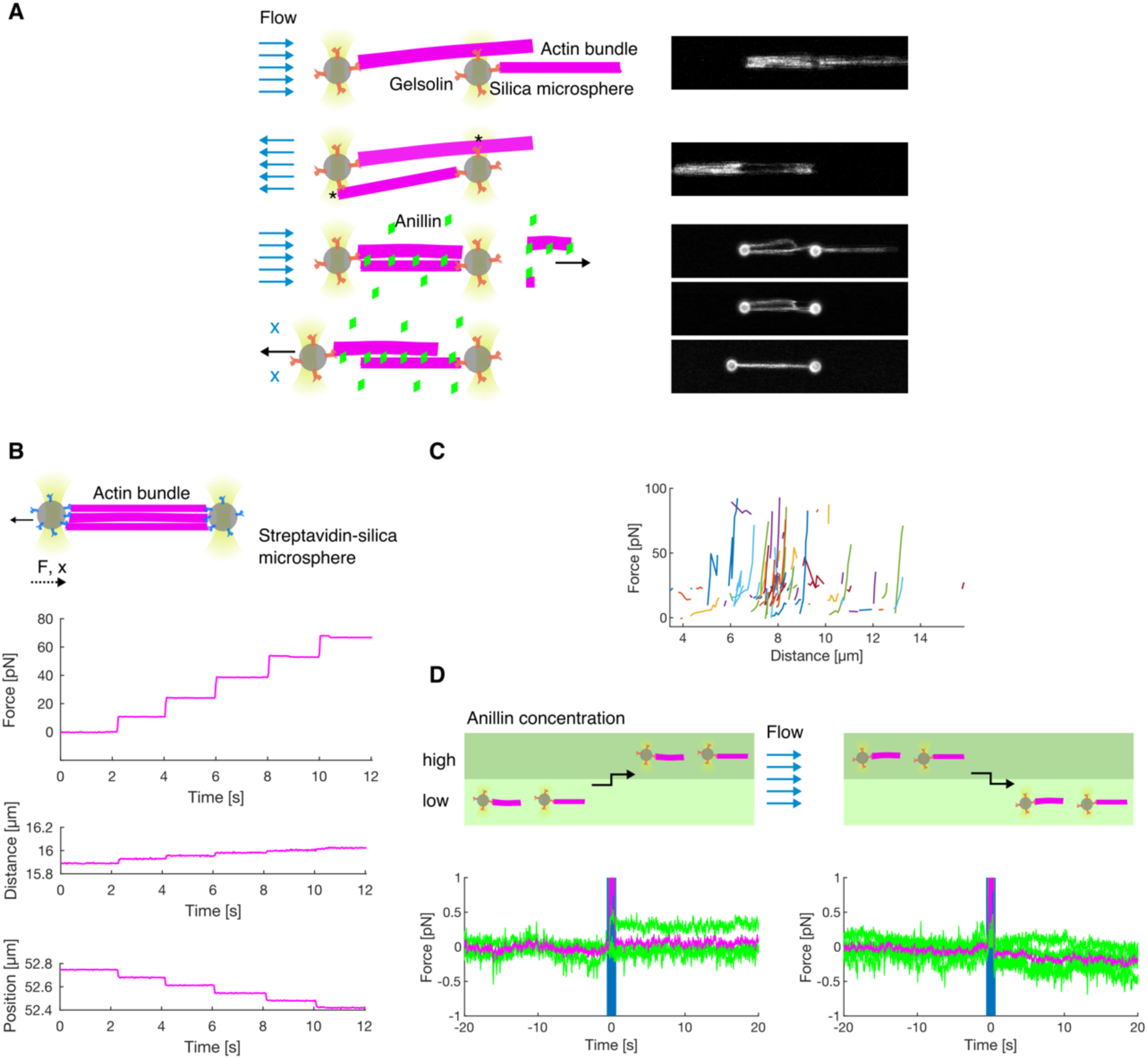
(**A**) Schematics and fluorescence micrographs showing how the anillin-actin bundle was created in the optical tweezers. The actin-anillin bundle used in the optical tweezers experiment was created similarly as in (*4*). Briefly, the barbered ends of the filaments firstly bind gelsolin-coated microspheres. Then, by reversing the buffer flow, the filaments are transiently attached (indicated by asterisk) to the opposite microsphere. The construct is then moved to anillin-GFP channel where the higher concentration of CaCl_2_ triggers severing activity of gelsolin. As anillin binds the filaments at the same time, its bundling activity prevents the construct from breaking up (Methods). (**B**) Control experiment showing the force time trace (top) and the distance between the beads (middle) in response to step-wise movement of the left microsphere (bottom) when the actin bundle was bridging both microspheres (biotin-phalloidin stabilised actin filaments suspended between two streptavidin microspheres). In contrast to Fig. 4 C, the force time trace increases in steps concomitant with the microsphere position steps. The steps in detected force are not followed by relaxation, showing that the actin filaments in the bundle form a solid bridge between the two microspheres and cannot slide relative to each other. (**C**) Steady state asymptotical force values of all measured force-distance curves of anillin-bundled actin filaments. Individual experiments are colour-coded (n = 43 actin-anillin bundles in 43 experiments). (**D**) Control experiments showing the force time trace when the actin-decorated but disjoined microspheres were moved between two channels with different concentration of anillin-GFP. Moving the disjoined construct between microfluidic channels with a stationary laminar flow (schematic depiction on the top) does not cause transient changes in the force response (bottom) - compare to Fig. 4 G. Green curves represent individual measurements (n = 8 events in 3 experiments) with mean shown in magenta.

**Movie S1.**

Diffusive motion of mobile actin filaments crosslinked by anillin (not visualized) to long, sparsely fluorescently labelled, immobilised actin filaments.

**Movie S2.**

Anillin-driven sliding of a mobile actin filament (bright) along an immobilised filament (dim).

**Movie S3.**

Anillin-driven sliding of two mobile actin filaments.

**Movie S4.**

Actin filaments do not bundle in the absence of anillin.

**Movie S5.**

Anillin couples with actin disassembly to generate directed filament sliding.

**Movie S6.**

Constriction of the anillin-actin ring composed of non-stabilised actin filaments. Overlay of the anillin-GFP (cyan) and rhodamine-actin (magenta) fluorescence channels.

## Materials and Methods

### Proteins

Homo sapiens anillin (GeneBank accession number: BC070066) cDNA was purchased (BC070066-seq-TCHS1003-GVO-TRI, BioCat GmbH Heidelberg, Germany), PCR amplified and ligated into an AscI-NotI-digested pOCC destination vector (*1*) containing a C-terminal GFP tag followed by a 3C PreScission Protease cleavage site and a 6xHis-tag. The protein was expressed in SF9 insect cells using an opensource FlexiBAC baculovirus vector system (*1*). The insect cells were harvested after 4 days by centrifugation at 300 g for 10 min at 4 °C in an Avanti J-26S ultracentrifuge (JLA-9.1000 rotor, Beckman Coulter). The cell pellet was resuspended in 5 ml ice-cold phosphate buffered saline (PBS) and stored at - 80 °C for further use. For cell lysis, the insect cells were homogenized in 30 ml ice-cold His-Trap buffer (50 mM Na-phosphate buffer, pH 7.5, 5% glycerol, 300 mM KCl, 1 mM MgCl_2_, 0.1% tween-20, 10 mM BME, 0.1 mM ATP) supplemented with 30 mM imidazole, Protease Inhibitor Cocktail (cOmplete, EDTA free, Roche) and benzonase to the final concentration of 25 units/ml, and centrifuged at 45000 x g for 60 min at 4 °C in the Avanti J-26S ultracentrifuge (JA-30.50Ti rotor, Beckman Coulter). The cleared cell lysate was incubated in a lysis buffer-equilibrated Ni-NTA column (HisPur Ni-NTA Superflow Agarose, Pierce, VWR) for 2 h at 4 °C on a rotator. The Ni-NTA column was washed with the wash buffer (His-Trap buffer supplemented with 60 mM imidazole) and the protein was eluted with the elution buffer (His-Trap buffer supplemented with 300 mM Imidazole). The fractions containing anillin-GFP were pooled, diluted 1:10 in the His-Trap buffer and the purification tag was cleaved overnight with 3C PreScisson protease. The solution was reloaded onto a Ni-NTA column to further separate the cleaved protein from the 6xHis-tag. The protein was concentrated using an Amicon ultracentrifuge filter and flash frozen in liquid nitrogen. Protein concentration was measured using Bradford assay (23236, Thermo Scientific) and microplate reader spectrometer (CLARIOstar, BMG Labtech, Germany). Kinesin-1 was expressed and purified as described previously (*2*).

Unlabelled rabbit muscle actin (AKL99, Cytoskeleton, Inc., USA) and rhodamine labelled rabbit muscle actin (AR05, Cytoskeleton, Inc., USA) was resuspended to final concentration of 10 mg/ml in general actin buffer (GAB; 5 mM TRIS-HCl pH 8, 0.2 mM CaCl_2_) supplemented with 0.2 mM ATP, 5% (w/v) sucrose and 1% (w/v) dextran. For experiments with fluorescently labelled non-stabilized filaments, rhodamine labelled actin was diluted with the unlabelled actin above in a 1:10 ratio. The actin was then aliquoted, flash-frozen and stored in −80 °C. Actin filaments were polymerised by mixing the aliquoted actin (0.32 mg/ml final concentration) with polymerisation buffer (5 mM Tris-HCl pH 8.0, 0.2 mM CaCl_2_, 50 mM KCl, 2 mM MgCl_2_, 1 mM ATP) and, optionally, phalloidin (15 µM final concentration) for filament stabilisation and additional labelling. Biotinylated phalloidin (biotin-XX phalloidin, B7474, Thermo Fisher Scientific), rhodamine-labelled phalloidin (R415, Thermo Fisher Scientific), or a 3:4 mixture of both were used in the experiments. Phalloidin-stabilised filaments were polymerised overnight at 4° C and remained stable for at least two months. Non-stabilised filaments were grown in the polymerisation solution in the absence of phalloidin for a period of 5 minutes to 5 hours and were used immediately after polymerization. For use in the assays, actin filaments were diluted in a ratio between 1:10 and 1:300, depending on the experiment.

Fluorescently-labelled microtubules were polymerized from 4 mg/ml porcine tubulin (80% unlabelled and 20% Alexa Fluor 647 NHS ester-labelled; Thermo Fisher Scientific) for 2 h at 37 °C in BRB80 (80 mM PIPES, 1 mM EGTA, 1 mM MgCl_2_, pH 6.9) supplemented with 1 mM MgCl_2_ and 1 mM GMPCPP (Jena Bioscience, Jena, Germany). The polymerized microtubules were centrifuged for 30 min at 18000 x g in a Microfuge 18 Centrifuge (Beckman Coulter, Brea, CA) and the pellet was resuspended in BRB80 supplemented with 10 µM taxol (BRB80T).

### Fluorescence imaging

The imaging was carried out in flow channels assembled from glass coverslips held together by parafilm spacers. Dichlorodimethylsilane-treated glass was passivated by 1% F127 pluronic copolymer (P2443, Sigma) for at least 30 minutes. Attachment of actin filaments stabilised by biotin-phalloidin was performed in channels that were incubated with an anti-biotin antibody solution (1 mg/ml in PBS, B3640, Sigma) prior to passivation. All channels were washed with 40 µl of the assay buffer upon passivation.

Actin filaments (either labelled by rhodamine or stabilised by rhodamine-phalloidin) and anillin-GFP were imaged using an inverted microscope (Ti-E Eclipse, Nikon) equipped with 60x and 100x 1.49 N.A. oil immersion objectives (CFI Apo TIRF and HP Apo TIRF, respectively, Nikon) in a TIRF (total internal refraction fluorescence) regime. The GFP and rhodamine fluorophores were sequentially excited by a laser on the wavelengths of 488 nm and 567 nm or 561nm, respectively. FITC and TRITC fluorescence filter cubes were used, and the fluorescence was recorded using a CCD camera (iXon Ultra DU888, Andor Technology) or a CMOS camera (sCMOS ORCA 4.0 V2, Hamamatsu Photonics). The imaging setup was controlled by NIS Elements software (Nikon). The frame rate ranged between 1 frame per 10 ms to 1 frame per minute - time is indicated as time scale bars in kymographs, or as timestamps in time lapse micrographs. The experiments were performed at room temperature.

### Anillin binding and diffusion assay

Actin filaments stabilised by biotin-phalloidin were attached to anti-biotin antibodies on a glass coverslip. The channel was then washed with twice the channel volume of HEPES-based imaging buffer (20 mM HEPES, 2 mM MgCl_2_, 1 mM EGTA, pH 7.2 KOH, 10 mM DTT, 20 mM D-glucose, 0.1 % Tween-20, 0.5 mg/ml Casein, 1 mM ATP, 0.22 mg/ml Glucose Oxidase, 0.02 mg/ml Catalase) before the HEPES-based imaging buffer with 0.12 nM anillin-GFP was flushed into the channel.

The oligomeric state of the anillin-GFP was determined from the fluorescence intensity of individual molecules diffusing on the biotin-immobilised actin filaments. This signal was compared with the fluorescence intensity of a single kinesin-1-GFP molecules immobilised on microtubules in the presence of AMP-PNP (in the absence of ATP). Kinesin-1-GFP was used as a standard due to its known dimeric state and therefore two-step photobleaching intensity profile (*3*). Biotin-labelled microtubules were immobilised in the flow channel similarly to the actin filaments. The concentration of the kinesin-1-GFP was adjusted such that individual molecules bound to microtubules were detectable.

Kinetic properties of anillin-GFP unbinding from actin filaments was estimated from the temporal profile of the fluorescence intensity integrated along the contour of immobilised actin filaments upon the introduction, or washout of anillin-GFP, to or from, the imaging chamber. This process was repeated with various concentrations (from 0.12 to 12 nM) of the anillin-GFP in order to obtain a concentration profile.

### Actin filament sliding assay

Biotin-phalloidin stabilised actin filaments were introduced in the flow channel as described above. The channel was subsequently washed with the imaging buffer. Then, non-biotinylated actin filaments diluted in the imaging buffer containing 12 nM anillin-GFP were flushed into the channel. These filaments formed bundles with the biotin-phalloidin-stabilised filaments attached to the glass surface.

In an alternative experiment, approx. 1 µl of actin filaments followed by 15 µl of the imaging buffer with 0.1% methylcellulose (0.1 % w/v final concentration, 4000 cps at 2 %, M0512, Sigma, supplemented with 30 mM NaCl final concentration) were first introduced into the flow channel. After initial imaging, the channel was flushed with imaging buffer containing 12 nM anillin-GFP and 0.1% methylcellulose. Filaments that remained in the channel after the flush were observed forming bundles.

### Actin filament disassembly assay

Latrunculin-driven disassembly experiments were performed by introducing long unlabelled non-stabilised actin filaments into the flow channel, followed by short rhodamine-phalloidin stabilised actin filaments and 12 nM anillin-GFP in the GAB imaging buffer (GAB buffer with 10 mM DTT, 20 mM D-glucose, 0.1 % Tween-20, 0.5 mg/ml Casein, 1 mM ATP, 0.8 mM PIPES, 0.22 mg/ml Glucose Oxidase, 0.02 mg/ml Catalase) supplemented with 0.1% methylcellulose and 80-200 nM latrunculin (L12370, Thermo Fisher Scientific).

### Actin ring constriction assay

For the ring constriction experiments, actin filaments were diluted 1:10 in the imaging buffer supplemented with 12 nM anillin-GFP and 0.1% methylcellulose. The final mixture was immediately transferred into the flow channel for imaging. We tested the robustness of the constriction under various buffer conditions. Ring constrictions were observed in both HEPES-based and GAB-based imaging buffers, with or without an additional supplement of 50 mM KCl. The data were pooled into two groups based on the properties of the actin filaments (phalloidin stabilised vs. non-stabilised) since the statistical testing (two-sample t-test at the significance level 0.05) did not reject the hypothesis that the constrictions rates within these groups come from distributions of equal means.

### Optical trapping assay

Correlative force measurements and microscopy were performed on an optical tweezers setup equipped with confocal fluorescence imaging and microfluidic system (c-Trap, LUMICKS, The Netherlands). The microspheres, actin filaments, imaging buffers and proteins were simultaneously flushed in the 4-channel air-pressure driven microfluidic system in a laminar flow mode, ensuring that the solutions did not mix. The microfluidic system was passivated by BSA (0.1% in PBS) and F127 Pluronic no later than 100 hours before the experiment. The trap stiffness was calibrated using the thermal spectrum method (Scanary v. 3.3.0, LUMICKS, The Netherlands) under zero flow condition. The experiments were performed at room temperature.

Actin filaments stabilised with rhodamine-phalloidin were attached to microspheres using barbered-end-binding protein gelsolin. Gelsolin-coated microspheres were prepared using a previously published protocol (*4*). Briefly, the 1.01 µm diameter carboxylated silica microspheres (SC04000, Bangs Beads) were activated by amine-reactive crosslinker chemistry using NHS (N-hydroxysuccinimide, 130672-5G, Sigma) and functionalized by gelsolin (HPG6-A, Cytoskeleton).

To form the actin bundles (Fig. S4 A), we modified a method described previously (*4*). Two microspheres were captured in two traps. These microspheres were moved to the actin channel (actin filaments stabilised by rhodamine-phalloidin in the GAB imaging buffer) where ends of the filaments were attached to both of the microspheres. These constructs were then moved to the channel with 2xPBS imaging buffer (274 mM NaCl, 5.4 mM KCl, 3.6 mM KH_2_HPO_4_, 16 mM Na_2_HPO_4_, 0.1 mM CaCl_2_, 10 mM DTT, 20 mM D-glucose, 0.1 % Tween-20, 0.5 mg/ml Casein, 1 mM ATP, 0.22 mg/ml Glucose Oxidase, 0.02 mg/ml Catalase) where we used a flow reversal for transient sideways binding of filaments to the opposite microsphere. This construct was immediately moved to the channel with 12 nM anillin-GFP in the GAB imaging buffer, where gelsolin-dependent severing of the sideway attached filaments occurred, and the anillin-GFP formed the actin bundle at the same moment. The flow in the channels was then stopped, and the force response of the construct to stretching and relaxation was measured as follows.

The microspheres were moved relative to each other in the direction of the bundle using the Trap Stepper utility of the c-trap software (v. 3.6.1, LUMICKS, The Netherlands), with 100 nm step size and 3 µm/s speed. The distance of the microspheres was measured using bright-field microscopy, and the visualisation of the actin filaments and anillin-GFP was provided by built-in scanning confocal microscope with 488 nm and 561 nm excitation lasers. The number of actin filaments in the bundle was estimated from the intensity of the fluorescence signal of the rhodamine-phalloidin. The acquisition was controlled by Scanary software (v. 3.3.0, LUMICKS, The Netherlands).

The response of the bundle to a change of the anillin-GFP concentration was measured in a 5-channel microfluidic system. Similarly as above, the actin-anillin bundle was formed and pre-stretched in 12 nM anillin-GFP channel. The construct was then moved to the channel with 1 nM anillin-GFP concentration. Due to the different geometry of the channels, the experiment was performed in the flow of about 1 µl/min to prevent mixing of the low and the high concentration anillin-GFP channels. Control experiments were performed to exclude the effects of the flow on the accuracy of the force measurement (Fig. S4 D).

### Image and data analysis

The oligomeric state of anillin-GFP and its diffusion behaviour were quantified using FIESTA (*5*) for the tracking of single molecules and @msdanalyzer MATLAB tool (*6*) for data analysis. The background signal was calculated from the areas directly adjacent to the actin filaments or microtubules and subtracted from the fluorescence intensities for quantification. The binding kinetics of anillin-GFP was estimated from the time constant of the first-order fit to the temporal step-response concentration profiles using a custom MATLAB (Mathworks) procedures. These profiles were measured as the anillin-GFP fluorescence intensity integrated along the contour of immobilised actin filaments upon the introduction or washout of anillin-GFP to, or from the flow chamber (see Fig. S1 A).

The lengths of the actin filament overlaps, as well as the circumferences of the rings, were measured manually in FiJi (*7*). The circumferences of the rings were measured through the centre of the contour. Kymographs were generated using MultiKymograph plugin in FiJi. The data were analysed by self-written MATLAB scripts. Exponential decays in increasing and decreasing forms, 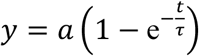, and 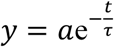, respectively, were used to fit transient events. Violin plots were generated using scripts by Bastian Bechtold. Raw images were treated with automatic brightness/contrast enhancement routine (as implemented in FiJi) for presentation purposes. Temporal profiles of the force and distance observed in the optical trapping assay were analysed in a self-written MATLAB procedure.

### Reproducibility and data exclusion

All replication attempts that did not suffer from technical problems (such as improper channel passivation or malfunction of the instruments) were successful. All experiments were independently repeated at least three times, if not stated otherwise. The number of experiments, i.e. the number of individually filled channels in the case of microscopy assays and the number of constructs assembled *de novo* in the case of optical trapping assay, is indicated in the text.

## References

1. T. E. Schroeder, Cytokinesis: Filaments in the cleavage furrow. Exp. Cell Res. 53, 272–318 (1968).

2. K. Fujiwara, T. D. Pollard, Fluorescent antibody localization of myosin in the cytoplasm, cleavage furrow, and mitotic spindle of human cells. J. Cell Biol. 71, 848–875 (1976).

3. T. D. Pollard, B. O’Shaughnessy, Annu. Rev. Biochem., 88, 12.1–12.29 (2019).

4. C. DeKraker, E. Boucher, C. A. Mandato, Regulation and assembly of actomyosin contractile rings in cytokinesis and cell repair. Anat. Rec. 301, 2051–2066 (2018).

5. M. C. Mangione, K. L. Gould, Molecular form and function of the cytokinetic ring. J. Cell Sci. 132, jcs226928 (2019).

6. H. E. Huxley, J. Hanson, Changes in the cross-striations of muscle during contraction and stretch and their structural interpretation. Nature. 173, 973–976 (1954).

7. A. F. Huxley, R. Niedergerke, Structural changes in muscle during contraction. Nature. 4412, 971–973 (1954).

8. T. Kamasaki, M. Osumi, I. Mabuchi, Three-dimensional arrangement of F-actin in the contractile ring of fission yeast. J. Cell Biol. 178, 765–771 (2007).

9. H. Ennomani, G. Letort, C. Guérin, J. L. Martiel, W. Cao, F. Nédélec, E. M. De La Cruz, M. Théry, L. Blanchoin, Architecture and Connectivity Govern Actin Network Contractility. Curr. Biol. 26, 616–626 (2016).

10. M. Murrell, P. W. Oakes, M. Lenz, M. L. Gardel, Forcing cells into shape: The mechanics of actomyosin contractility. Nat. Rev. Mol. Cell Biol. 16, 486–498 (2015).

11. T. H. Cheffings, N. J. Burroughs, M. K. Balasubramanian, Actomyosin Ring Formation and Tension Generation in Eukaryotic Cytokinesis. Curr. Biol. 26, R719–R737 (2016).

12. X. Ma, M. Kovacs, M. A. Conti, A. Wang, Y. Zhang, J. R. Sellers, R. S. Adelstein, Nonmuscle myosin II exerts tension but does not translocate actin in vertebrate cytokinesis. Proc. Natl. Acad. Sci. 109, 4509–4514 (2012).

13. E. Bi, P. Maddox, D. J. Lew, E. D. Salmon, J. N. McMillan, E. Yeh, J. R. Pringle, Involvement of an actomyosin contractile ring in Saccharomyces cerevisiae cytokinesis. J. Cell Biol. 142, 1301–1312 (1998).

14. R. Neujahr, C. Heizer, G. Gerisch, Myosin II-independent processes in mitotic cells of Dictyostelium discoideum: Redistribution of the nuclei, re-arrangement of the actin system and formation of the cleavage furrow. J. Cell Sci. 110, 123–137 (1997).

15. Z. Xue, A. M. Sokac, Back-to-back mechanisms drive actomyosin ring closure during Drosophila embryo cleavage. J. Cell Biol. 215, 335–344 (2016).

16. C. P. Descovich, D. B. Cortes, S. Ryan, J. Nash, L. Zhang, P. S. Maddox, F. Nedelec, A. S. Maddox, Crosslinkers both drive and brake cytoskeletal remodeling and furrowing in cytokinesis. Mol. Biol. Cell, 150813 (2018).

17. P. Bun, S. Dmitrieff, J. M. Belmonte, F. J. Nédélec, P. Lénárt, A disassembly-driven mechanism explains F-actin-mediated chromosome transport in starfish oocytes. Elife. 7, 1–27 (2018).

18. D. B. Oelz, B. Y. Rubinstein, A. Mogilner, Article A Combination of Actin Treadmilling and Cross-Linking Drives Contraction of Random Actomyosin Arrays. Biophys. J. 109, 1818–1829 (2015).

19. S. X. Sun, S. Walcott, C. W. Wolgemuth, Cytoskeletal cross-linking and bundling in motor-independent contraction. Curr. Biol. 20, R649–R654 (2010).

20. I. Mendes Pinto, B. Rubinstein, A. Kucharavy, J. R. Unruh, R. Li, Actin Depolymerization Drives Actomyosin Ring Contraction during Budding Yeast Cytokinesis. Dev. Cell. 22, 1247–1260 (2012).

21. M. Braun, Z. Lansky, A. Szuba, F. W. Schwarz, A. Mitra, M. Gao, A. Lüdecke, P. R. Ten Wolde, S. Diez, Changes in microtubule overlap length regulate kinesin-14-driven microtubule sliding. Nat. Chem. Biol. 13, 1245–1252 (2017).

22. Z. Lansky, M. Braun, A. Luedecke, M. Schlierf, P. R. Ten Wolde, M. E. Janson, S. Diez, Diffusible Crosslinkers Generate Directed Forces in Microtubule Networks. Cell. 160, 1159–1168 (2015).

23. B. Akiyoshi, K. K. Sarangapani, A. F. Powers, C. R. Nelson, S. L. Reichow, H. Arellano-Santoyo, T. Gonen, J. A. Ranish, C. L. Asbury, S. Biggins, Tension directly stabilizes reconstituted kinetochore-microtubule attachments. Nature. 468, 576–579 (2010).

24. M. Braun, Z. Lansky, F. Hilitski, Z. Dogic, S. Diez, Entropic forces drive contraction of cytoskeletal networks. BioEssays. 38, 474–481 (2016).

25. L. Zhang, A. S. Maddox, Anillin. Curr. Biol. 20, 135–136 (2010).

26. C. M. Field, B. M. Alberts, Anillin, a contractile ring protein that cycles from the nucleus to the cell cortex. J. Cell Biol. 131, 165–178 (1995).

27. P. Paolo D’Avino, How to scaffold the contractile ring for a safe cytokinesis - lessons from Anillin-related proteins. J. Cell Sci. 122, 1071–1079 (2009).

28. H. Kueh, W. Brieher, T. Mitchison, Dynamic stabilization of actin filaments. Proc. Natl. Acad. Sci. USA. 105, 16531 (2008).

29. M. Mavrakis, Y. Azou-Gros, F. C. Tsai, J. Alvarado, A. Bertin, F. Iv, A. Kress, S. Brasselet, G. H. Koenderink, T. Lecuit, Septins promote F-actin ring formation by crosslinking actin filaments into curved bundles. Nat. Cell Biol. 16, 322–334 (2014).

30. A. W. C. Lau, A. Prasad, Z. Dogic, Condensation of isolated semi-flexible filaments driven by depletion interactions. Europhys. Lett. 87, 48006 (2009).

31. A. Ward, F. Hilitski, W. Schwenger, D. Welch, A. W. C. Lau, V. Vitelli, L. Mahadevan, Z. Dogic, Solid friction between soft filaments. Nat. Mater. 14, 583–588 (2015).

32. F. Hilitski, A. R. Ward, L. Cajamarca, M. F. Hagan, G. M. Grason, Z. Dogic, Measuring cohesion between macromolecular filaments one pair at a time: Depletion-induced microtubule bundling. Phys. Rev. Lett. 114, 1–6 (2015).

33. T. Sanchez, I. M. Kulic, Z. Dogic, Circularization, photomechanical switching, and a supercoiling transition of actin filaments. Phys. Rev. Lett. 104, 65–68 (2010).

34. D. J. Odde, Previews and the Ideal Gas Law. Cell. 160, 1041–1043 (2015).

35. V. Bormuth, V. Varga, J. Howard, E. Schäffer, Protein Friction Limits Diffusive and Directed Movements of Kinesin Motors on Microtubules. Science. 325, 870–873 (2007).

36. F. Gittes, B. Mickey, J. Nettleton, J. Howard, Flexural rigidity of microtubules and actin filaments measured from thermal fluctuations in shape. J. Cell Biol. 120, 923–934 (1993).

37. M. F. Norstrom, P. A. Smithback, R. S. Rock, Unconventional processive mechanics of non-muscle myosin IIB. J. Biol. Chem. 285, 26326–26334 (2010).

38. M. R. Stachowiak, C. Laplante, H. F. Chin, B. Guirao, E. Karatekin, T. D. Pollard, B. O’Shaughnessy, Mechanism of cytokinetic contractile ring constriction in fission yeast. Dev. Cell. 29, 547–561 (2014).

39. M. Lenz, T. Thoresen, M. L. Gardel, A. R. Dinner, Contractile units in disordered actomyosin bundles arise from f-actin buckling. Phys. Rev. Lett. 108, 1–5 (2012).

40. M. P. Murrell, M. L. Gardel, F-actin buckling coordinates contractility and severing in a biomimetic actomyosin cortex. Proc. Natl. Acad. Sci. 109, 20820–20825 (2012).

41. D. Vavylonis, J. Q. Wu, S. Hao, B. O’Shaughnessy, T. D. Pollard, Assembly mechanism of the contractile ring for cytokinesis by fission yeast. Science (80-.). 319, 97–100 (2008).

42. K. Kruse, F. Jülicher, Actively contracting bundles of polar filaments. Phys. Rev. Lett. 85, 1778–1781 (2000).

43. T. B. Liverpool, M. C. Marchetti, Bridging the microscopic and the hydrodynamic in active filament solutions. Europhys. Lett. 69, 846–852 (2005).

44. M. Mishra, J. Kashiwazaki, T. Takagi, R. Srinivasan, Y. Huang, M. K. Balasubramanian, I. Mabuchi, In vitro contraction of cytokinetic ring depends on myosin II but not on actin dynamics. Nat. Cell Biol. 15, 853–859 (2013).

45. N. Kaji, K. Ohashi, M. Shuin, R. Niwa, T. Uemura, K. Mizuno, Cell cycle-associated changes in Slingshot phosphatase activity and roles in cytokinesis in animal cells. J. Biol. Chem. 278, 33450–33455 (2003).

46. K. Ono, M. Parast, C. Alberico, G. M. Benian, S. Ono, Specific requirement for two ADF/cofilin isoforms in distinct actin-dependent processes in Caenorhabditis elegans. J. Cell Sci. 116, 2073–2085 (2003).

47. M. P. Somma, B. Fasulo, G. Cenci, E. Cundari, M. Gatti, Molecular Dissection of Cytokinesis by RNA Interference in Drosophila Cultured Cells. Mol. Biol. Cell. 13, 2448–2460 (2002).

48. D. R. Gestaut, B. Graczyk, J. Cooper, P. O. Widlund, A. Zelter, L. Wordeman, C. L. Asbury, T. N. Davis, Phosphoregulation and depolymerization-driven movement of the Dam1 complex do not require ring formation. Nat. Cell Biol. 10, 407–414 (2008).

49. M. K. Gardner, D. J. Odde, Dam1 complexes go it alone on disassembling microtubules. Nat. Cell Biol. 10, 379–381 (2008).

50. T. L. Hill, Theoretical problems related to the attachment of microtubules to kinetochores. Proc. Natl. Acad. Sci. U. S. A. 82, 4404–4408 (1985).

51. T. R. Arnold, J. H. Shawky, R. E. Stephenson, K. M. Dinshaw, T. Higashi, F. Huq, L. A. Davidson, A. L. Miller, Anillin regulates epithelial cell mechanics by structuring the medial-apical actomyosin network. Elife (2019), doi: 10.7554/eLife.39065.

## Supplementary References

1. R. P. Lemaitre, A. Bogdanova, B. Borgonovo, J. B. Woodruff, D. N. Drechsel, FlexiBAC: A versatile, open-source baculovirus vector system for protein expression, secretion, and proteolytic processing. BMC Biotechnol. 19, 1–11 (2019).

2. B. Nitzsche, V. Bormuth, C. Bräuer, J. Howard, L. Ionov, J. Kerssemakers, T. Korten, C. Leduc, F. Ruhnow, S. Diez, Studying kinesin motors by optical 3D-nanometry in gliding motility assays. Methods Cell Biol. 95, 247–271 (2010).

3. N. Hirokawa, K. K. Pfister, H. Yorifuji, M. C. Wagner, S. T. Brady, G. S. Bloom, Submolecular domains of bovine brain kinesin identified by electron microscopy and monoclonal antibody decoration. Cell. 56, 867–878 (1989).

4. A. Ward, F. Hilitski, W. Schwenger, D. Welch, A. W. C. Lau, V. Vitelli, L. Mahadevan, Z. Dogic, Solid friction between soft filaments. Nat. Mater. 14, 583–588 (2015).

5. F. Ruhnow, D. Zwicker, S. Diez, Tracking single particles and elongated filaments with nanometer precision. Biophys. J. 100, 2820–2828 (2011).

6. N. Tarantino, J. Y. Tinevez, E. F. Crowell, B. Boisson, R. Henriques, M. Mhlanga, F. Agou, A. Israël, E. Laplantine, Tnf and il-1 exhibit distinct ubiquitin requirements for inducing NEMO-IKK supramolecular structures. J. Cell Biol. 204, 231–245 (2014).

7. J. Schindelin, I. Arganda-Carreras, E. Frise, V. Kaynig, M. Longair, T. Pietzsch, S. Preibisch, C. Rueden, S. Saalfeld, B. Schmid, J. Y. Tinevez, D. J. White, V. Hartenstein, K. Eliceiri, P. Tomancak, A. Cardona, Fiji: An open-source platform for biological-image analysis. Nat. Methods. 9, 676–682 (2012).

8. A. W. C. Lau, A. Prasad, Z. Dogic, Condensation of isolated semi-flexible filaments driven by depletion interactions. EPL (Europhysics Lett. 87, 48006 (2009).

9. K. Matsuda, M. Sugawa, M. Yamagishi, N. Kodera, J. Yajima, Visualizing dynamic actin cross-linking processes driven by the actin-binding protein anillin. FEBS Lett. in press (2019), doi: 10.1002/1873-3468.13720.

